# Templating of Monomeric Alpha-Synuclein Induces Inflammation and SNpc Dopamine Neuron Death in a Genetic Mouse Model of Synucleinopathy

**DOI:** 10.1101/2024.07.29.605647

**Authors:** Matthew D. Byrne, Peyman Petramfar, Richard J Smeyne

**Author notes:** Corresponding author. Richard Jay Smeyne, Ph.D. Department of Neuroscience, Jefferson Comprehensive Parkinson’s Disease Center Thomas Jefferson University, 900 Walnut Street, Philadelphia PA 19107.

## Abstract

While the etiology of most cases of Parkinson’s disease (PD) are idiopathic, it has been estimated that 5-10% of PD arise from known genetic mutations. The first mutations described that leads to the development of an autosomal dominant form of PD are in the SNCA gene that codes for the protein alpha-synuclein (α-syn). α-syn is an abundant presynaptic protein that is natively disordered and whose function is still unclear. In PD, α-syn misfolds into multimeric β-pleated sheets that aggregate in neurons (Lewy Bodies/neurites) and spread throughout the neuraxis in a pattern that aligns with disease progression. Here, using IHC, HPLC, and cytokine analysis, we examined the sequelae of intraparenchymal brain seeding of oligomeric pre-formed fibrils (PFFs) and monomeric α-syn in C57BL/6J (WT) and A53T SNCA mutant mice. We found that injection of PFFs, but not monomeric α-syn, into the striatum of C57BL/6J mice induced spread of aggregated α-syn, loss of SNpc DA neurons and increased neuroinflammation However, in A53T SNCA mice, we found that both PFFs and monomeric α-syn induced this pathology. This suggests that the conformation changes in α-syn seen in the A53T strain can recruit wild-type α-syn to a pathological misfolded conformation which may provide a mechanism for the induction of PD in humans with SNCA duplication/triplication.

## Templating of Monomeric Alpha-Synuclein Induces Inflammation and SNpc Dopamine Neuron Death in a Genetic Mouse Model of Synucleinopathy

Parkinson’s disease (PD) is a multisystem neurodegenerative disorder affecting over six million people worldwide [1]. While the majority of PD cases are considered to be idiopathic, a number of well characterized genetic mutations have been identified that account for approximately 5-10% of those diagnosed with PD. Interestingly, the presentation of genetic and sporadic cases are often indistinguishable [2], including both the characteristic motor and non-motor symptoms. The classic motor symptoms of PD include bradykinesia, tremor, and rigidity, while the non-motor symptoms include gut dysfunction, sleep disorders, speech pathologies and even psychiatric issues such as depression [3, 4]. The onset of both the motor and non-motor symptoms is often insidious, but symptoms are progressive. At this time, there are no disease-modifying therapies available for patients [5]. In the CNS, PD pathology is characterized by the progressive loss of the dopaminergic (DAergic) neurons located in the substantia nigra pars compacta (SNpc), degeneration of terminals of these neurons leading to deficits in striatal dopamine, and neuroinflammation that includes astrocytosis, microgliosis and increased pro-inflammatory cytokine levels [6, 7]. However, the most characteristic feature of PD pathology is the presence of dense proteinaceous intraneuronal inclusions, called Lewy bodies, within neurons in the brain. Although many proteins are present in these inclusions, the most abundant protein is alpha synuclein (α-Syn) [8]. α-Syn was first linked to PD by the discovery of a kindred family who displayed an accelerated onset of PD symptoms, oft times at significantly earlier ages that the traditional age of PD symptom onset, with a dominant pattern of inheritance [9]. Subsequent studies have shown discovered families containing point mutations, duplications, and triplications of the SNCA gene that encodes α-Syn; all of which are causative for PD. Additional work has demonstrated an association between α-Syn levels and the risk of developing sporadic PD [10].

While the normal function of α-Syn is still poorly understood, it is an 140 amino acid protein that is enriched at the presynaptic terminal, and accounts for as much as 1% of all proteins in the cytosol of cells in the CNS [11]. Under homeostatic conditions, α-Syn exists as a disordered monomer, however, it is prone to aggregating into higher order soluble oligomeric species as well as insoluble aggregates, characterized by β-pleated sheets and phosphorylation at serine129 [11]. These insoluble aggregates are toxic *in vitro* and *in vivo*, however, their toxicity can depend on both the conformational and molecular weight of the assembly [12].

While the pathological hallmark of PD is the presence of Lewy bodies in the SNpc DA neurons, there is a predictable progression of α-Syn pathology, characterized by the progressive appearance of aggregated misfolded α-Syn in the neuroaxis that coincides with the clinical progression of the disease [13]. Temporally, this Lewy pathology first appears in the peripheral neurons of the gut followed by appearance in the dorsal motor nucleus of the vagus nerve (DMV) and olfactory bulb. Later, pathology is noted in the lower brainstem nuclei, the SNpc, and then finally, in some cases, to neocortical nuclei pathological burden [13]. The pattern of α-Syn misfolding, along direct circuits, suggests that α-Syn pathology is transmissible, This observation was further confirmed from studies that found that transplanted fetal DA neurons, when implanted into the brains of PD patients developed this Lewy pathology despite a significant age difference in the neurons [14].

While it is hypothesized that the spread and accumulation of α-Syn aggregates in neurons drives deleterious neurodegeneration in PD, recent studies have suggested that there is a significant contribution from neuroinflammation that can both create and/or trigger a pathogenic cellular environment that is permissive for α-Syn aggregation [15]. For example, it has been shown that α-Syn activates microglia which can synergize neurotoxicity in vitro [16]. It has also been shown that this process can initiates and perpetuate a toxic cycle where continued release of aggregated α-Syn from damaged neurons, induces activation of microglia and astrocytes which then leads to subsequent neurodegeneration [16]. While this link between neuronal cell death, α-Syn aggregation, and glial activation has been studied in PD, there is still much debate about the initiating event(s) of PD pathology and role of inflammation.

Since α-Syn has been confirmed to play a central pathogenic role in a number of well-known synucleinopathies, powerful models of disease have been generated that allow for the experimental interrogation of the mechanisms of synuclein pathogenesis. These include the development of several mouse models that contain known pathogenic mutations (A53T, A30P) in the SNCA gene. However, for the most part, none of these mouse models have been able to faithfully recreate the nigrostriatal degeneration and α-Syn aggregation present in humans with PD [17]. To overcome this issue, an in vivo model for PD was developed that specifically focused on the role of misfolded, aggregated α-Syn. Here, monomeric recombinant human α-Syn is incubated to generate insoluble aggregated preformed fibrils (PFFs). These PFFs can be disrupted by sonication into short, aggregated oligomers that are small enough to be taken into neurons [18]. Using this model, a single intracerebral inoculation with α-Syn PFF’s has been shown to induce a pathology that approximates that seen in PD including insoluble aggregates of α-Syn in neurons, synaptic dysfunction, dysregulation of striatal dopamine release, and even nigrostriatal degeneration.[19, 20]. The mechanism for this appears to be that these PFFs act as a “seed”, capable of triggering aggregation of non-misfolded endogenous α-Syn in cells [21].

In this study we examined the neuropathological consequences of intracerebral inoculation with α-Syn PFFs or α-Syn monomers injected in C57BL/6J wildtype mice or C57BL/6J mice harboring a double PAC transgene overexpressing the human A53T mutation on a mouse SNCA-/- background [22]. We hypothesized that the transgenic line overexpressing α-Syn would result in accelerated pathology compared to wild type animals. We found that injection of PFFs into the striatum or hippocampus of both C57BL/6J WT and A53T transgenic mice resulted in a predictable and progressive pattern of α-Syn pathology that was circuit specific. We found that injection of the monomeric forms of α-Syn into C57BL/6J mice did not induce any progressive synuclein pathology. However, when the monomeric form of α-Syn was intracerebrally injected into mice harboring the human A53T mutation, we observed the induction of a synucleinopathy that was indistinguishable from that seen following injection of PFFs. This suggests that dysregulated overexpression of α-Syn, when in sufficient quantity, can induce misfolding of normal synuclein. We hypothesize that this may be the reason why humans carrying the A53T or even overexpression (duplication or triplication of the wild-type SNCA gene) develop PD pathology at a significantly earlier time than seen in idiopathic or even other familial forms of Parkinson’s disease.

## Materials and Methods

### Animals

All animals used in this study were purchased from the Jackson Labs (Bar Harbor ME). C57BL/6J mice (Cat # 00064) or FVB;129S6-Tg(SNCA*A53T)1Nbm *Snca^tm1Nbm^* Tg(SNCA*A53T)2Nbm/J (Also known as: dbl-PAC-tg(SNCAA53T);Snca-/-) (A53T, Cat 010799) then backcrossed to C57BL/6J for >10 generations. It is also important to note that the transgenic mice are on a mouse SNCA-/- background, meaning that all synuclein expressed in these animals are the human species. All mice were 6-12 weeks old when PFFs were intracerebrally injected.

### Preparation of α-Syn PFFs and Quality Control

Recombinant human α-Syn expressed in E. Coli was obtained from Proteos (Kalamazoo, MI). The monomeric form of α-Syn was centrifuged at 4°C for 10 minutes at 15,000 x g. After centrifugation, the supernatant was removed and its protein concentration was and determined using a Nanodrop Model 2000 spectrophotometer (Fisher Scientific). For each sample the monomer was diluted to a final concentration of 5mg/ml in ~100mM NaCl, ~7.5mM Tris, and ~10mM phosphate and adjusted to pH 7.4. A 100ug aliquot was placed into an orbital shaker at 37°C for 7 days at 1,000 RPM to induce fibrillization. Successful fibrillization in confirmed via transmission electron microscopy and sedimentation assay, with greater amounts of protein in the pellet versus supernatant fraction (Suppl Fig 1).

### Preparation of Surgical Material

PFFs or monomeric α-Syn were diluted to 2ug/ul in sterile dPBS. These samples were then sonicated using a microtip sonicator at power level 2 for .5 second ON/.5 second OFF pulses, with pausing every 10 seconds to prevent excess heat and frothing. Following sonication, successful disruption of the fibrils was confirmed using transmission electron microscopy, with average fibril being ~50nm min length (Fig 1). Monomeric protein was centrifuged at 15,000xg at 4C before surgical session, with the supernatant retained for use in the surgical injections. Injections.

**Fig 1.**
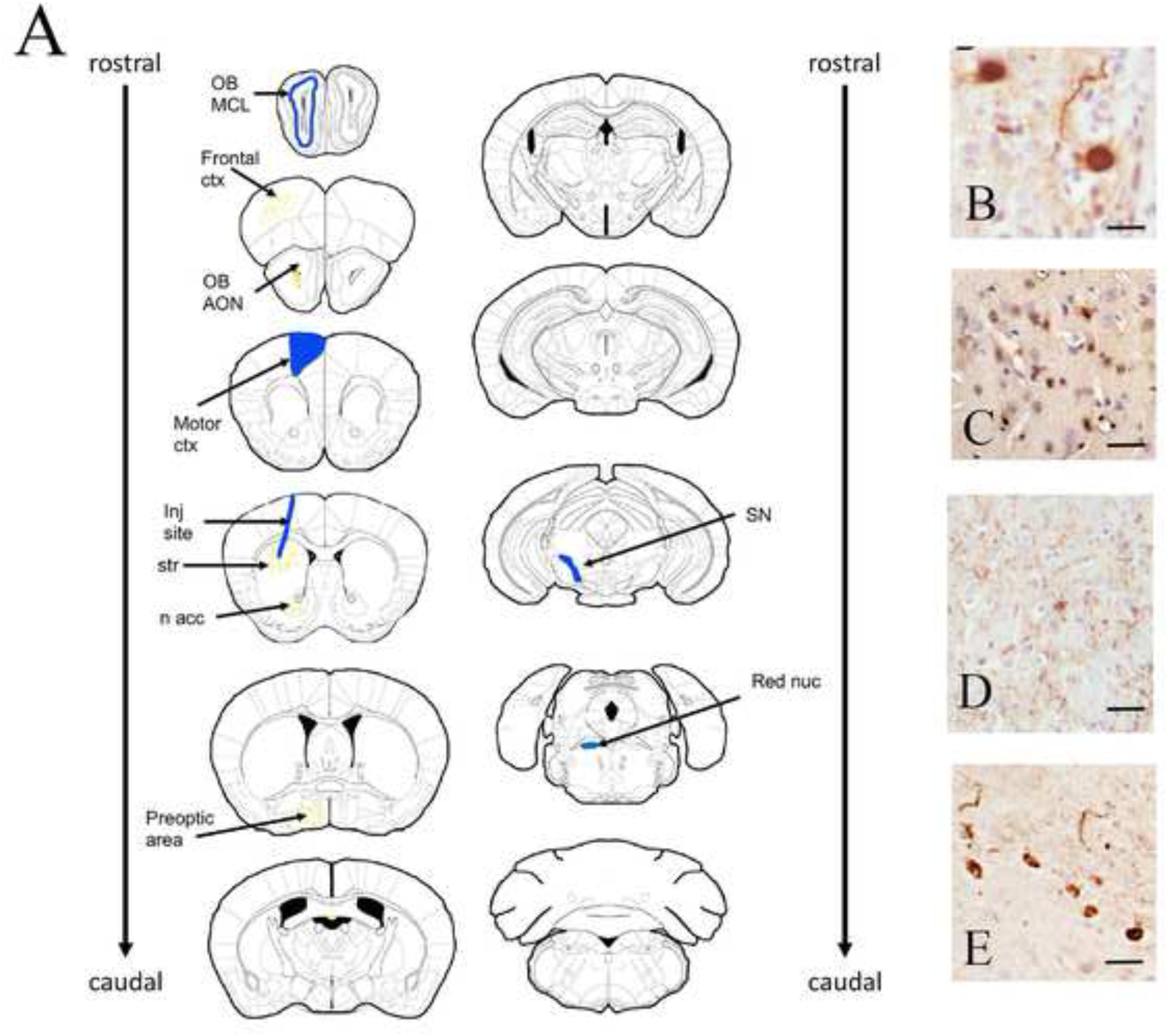
Map of pSer-129 aggregation in C57BL/6J and A53T mice following intrastriatal injection of PFFs. A. 60 days after intrastriatal injection of PFFs into the dorsolateral striatum we observed (from rostral to caudal) pSer129 α-syn immunoreactive aggregation throughout the neuraxis. Areas highlighted in blue showed medium to high levels of pSer129 α-syn aggregates, while areas in yellow showed low to medium aggregation. B. photomicrograph of pSer129 α-syn aggregate in the AON. C) photomicrograph of pSer129 α-syn aggregate in the motor cortex, D. photomicrograph of pSer129 α-syn aggregate in the striatum around the injection site, E. photomicrograph of pSer129 α-syn aggregate in the SNpc. OB: olfactory bub, MCL: mitral cell layer of the olfactory bulb, AON: accessory olfactory nucleus, inj site: injection site, str: striatum, n. acc: nucleus accumbens, SN: substantia nigra pars compacta. Scale bar B= 10 um, C-E, 25 um.

### Stereotaxic Injection of PFFS and monomeric α-Syn into Mouse Brain

Mice were anesthesized using a 3% mixture of isoflurane/oxygen. Following exposure of the skull, a small hole was drilled using a robotic stereotaxic instrument (Neurostar, Tubingen, Germany) at locations above the striatum or hippocampus. 2ul of diluted monomer or PFFs (2ug/ul) were injected into either the dorsal lateral striatum (AP 0.86, DV −2.5, ML 1.8) or rostral hippocampus (AP −2.18, DV −1.25, ML 1.75) using a 10ul Hamilton microsyringe at a constant rate of 0.4ul per minute. The syringe was left in place for 5 minutes after which it was slowly withdrawn. The scalp was then closed using proline suture and the wound was treated with topical lidocaine.

### Immunohistochemistry

30, 60, or 180 days post injection, mice were deeply anesthetizing with an intraperitoneal injection of 0.9% Avertin. After loss of corneal and deep tendon reflexes, mice were transcardially perfused with phosphate-buffered saline (PBS, pH 7.4), followed by 3% paraformaldehyde in PBS, pH 7.4. Brains were then dissected out of the skull following perfusion and post-fixed overnight in fresh 3% PFA. Brains were then dehydrated using a gradient solution of ethanol, defatted in xylenes, and then embedded in paraffin (Paraplas-Xtra, Fisher Scientific), and cut serially at 10um. Every section from the olfactory bulb to the anterior aspects of the cerebellar-midbrain junction was mounted to Superfrost-Plus slides (Fisher Scientific), five sections per slide. Every fourth slide (first series) was immunostained for p129-synuclein (1:40k, mouse monoclonal, 81a ab184674, abcam Cambridge, UK). The next slide in the series (second series) was immunolabeled for both tyrosine hydroxylase (TH) as a marker of dopaminergic neurons (1:250; mouse monoclonal, T1299, Sigma-Aldrich, MO, USA) and ionized calcium-binding adapter molecule 1 (Iba-1) (1:200; rabbit polyclonal, 019-19741, Wako, VA, USA). The double labeling was carried out using a two-color DAB/VIP protocol. All sections were counterstained with a Nissl stain (cresyl violet).

### Qualitative Determination of p129-synuclein burden

Sections from all brains in the anatomical studies, spaced 200 um apart were immunostained for pSer129 α-syn. Sections were examined under 20x objectives for the presence of insoluble aggregates of pSer129 α-syn. Each region identified in **Suppl Table 1** was examined and scored by an observer blinded to the experimental condition as being having low (0-20% of cells in the region containing pSer129 α-syn aggregates, medium (20-50% of cells in the region containing pSer129 α-syn aggregates) or high (greater than 50% of cells in the region containing pSer129 α-syn aggregates) based on the highest expression on a section within that region. Once the scoring for each animal was complete, the animals were unblinded and the relative SYN burden was determined. This was done by examining the scores for each structure in each animal in the group. Animals scored as low were assigned an ordinate score of 0, medium were assigned an ordinate score of 2 and those scored as high were assigned an ordinate score of 4. For each group, the ordinate numbers were summed and divided by the total number of animals. This average number was used to determine a relative burden for each structure in each experimental group.

### Stereological Assessment of SNpc DA neurons and microglia

The total number of SNpc TH-positive DA neurons (+ Nissl) were estimated by model-based stereology [23] using the physical disector (StereoInvetigator, MBF Bioscience). DA neurons were estimated from the entire rostral to caudal aspect of the SN. Microglia numbers were stereologically estimated using design-based stereology (optical dissector, StereoInvestigator, MBF Biosciences) [24]. Microglia were deemed as “resting” if the Iba-1-positive cell body was <3 microns in diameter and also possessed long slender processes. Microglia were counted as activated when the cell body was >3 microns but also had an irregular shape with shorter and thickened processes [25].

### Determination of striatal catecholamine content

C57BL/6J mice were deeply anesthetized with Avertin and transcardially perfused with 0.9% saline to remove the majority of the blood from the brain vasculature. Brains were rapidly removed and placed in a brain matrix (Model BS-AL-5000C, Braintree Scientific, Braintree. MA) and sliced into 1 mm thick sections. Dissected sections were placed on an ice-cooled plate and tissue was dissected from the SN (Bregma: −2.70 - −3.70), striatum (Bregma: +0.14-+1.26mm), brainstem (Bregma: −5.40 - −6.70mm), cortex (Bregma: −1.70 - −2.70mm) and the hippocampus (Bregma: −1.70 - −2.70mm) were dissected [26] and rapidly frozen on dry ice until processed. Tissues were homogenized, using a handheld sonic tissue dismembrator, in 100-750 ul of 0.1M TCA containing 0.01M sodium acetate, 0.1mM EDTA, and 10.5 % methanol (pH 3.8). Ten microliters of this homogenate was used for the protein assay. The samples were then spun in a microcentrifuge at 10,000 g for 20 minutes. Supernatant was removed for HPLC-ECD analysis. HPLC was performed using a Kinetix 2.6um C18 column (4.6 x 100 mm, Phenomenex, Torrance, CA USA). The same buffer used for tissue homogenization is used as the HPLC mobile phase.

For final determination of catecholamine content, the protein concentration in cell pellets was determined by BCA Protein Assay Kit (Thermo Scientific). Ten microliter tissue homogenates were distributed into 96-well plate and 200 ml of mixed BCA reagent (25 ml of Protein Reagent A is mixed with 500 μl of Protein Reagent B) was added. Plates were incubated at room temperature for two hours for the color development. A BSA standard curve is run at the same time. Absorbance is measured by the plate reader (POLARstar Omega), purchased from BMG LABTECH Company.

### Cytokine Assay

To examine the expression of cytokine/chemokines, brain tissue, prepared as described for HPLC analysis, was mechanically homogenized in Bioplex cell lysis buffer containing factors 1 and 2 (Bio-Rad, CA, USA) and centrifuged at 4500xg. The total protein concentration of each sample was determined using the BCA assay (Pierce, IL, USA), with bovine serum albumin as a standard, per the manufacturer’s protocol. Individual vials of cytokine/chemokine beads were sonicated, vortexed and then mixed with assay buffer (Milliplex Map Kit, MCYTOMAG-70K, Millipore, MA, USA). Working standards were made by diluting the stock concentration (10,000 pg/ml) in assay buffer. Brain and serum samples were added in equal volumes (25ul) into the wells of a 96-well plate containing 25ul of assay buffer and 25ul of cytokines mixed beads and plate was incubated overnight at +4C. Then the plate was washed and incubated with the following: (i) detection antibodies, (ii) Straptavidin-Phycoerythrin, (iii) sheath fluid. The plates were then run on a Luminex 200^TM^ according to the manufacturer’s recommended procedures and the data analyzed using BioPlex Manager 4.1 software (Bio-Rad, Hercules, CA). All samples were run in duplicates with data expressed as pg/mg total protein.

## Results

### Propagation of a-syn pathology after intracerebral injection of α-syn

To determine the propagation of α-syn pathology in the mouse brain following intracerebral α-syn injections, we injected human α-syn PFFs or monomer into either hippocampus or striatum, in both wild type C57Bl/6J and A53T mice. At 30, 60, or 180 days post injection, animals were transcardially perfused with 3% paraformaldehyde and brains were collected, serially sectioned and immunostained for the presence of non-soluble pSer129 α-syn. Immunostained sections throughout the whole of the neuraxis were graded for pSer129α-syn using a validated qualitative rating scale [27, 28] (supplement table 1); with scoring based on each region’s highest synuclein burden. Following injections of PFF’s into the dorsal striatum of both C57BL/6J and A53T animals, we observed a distribution of pSer129 α-syn throughout the structures of the nigrostriatal pathway (**Fig 1**). In C57BL/6J animals, pSer129 α-syn was first seen outside of the striatum at 30 days post injection and the expression was maximal at 60dpi. When pSer129 α-syn expression was examined at 180 dpi, we observed qualitatively lower levels of these insoluble aggregates than were seen at 60 days (**Fig 2**).

**Fig 2.**
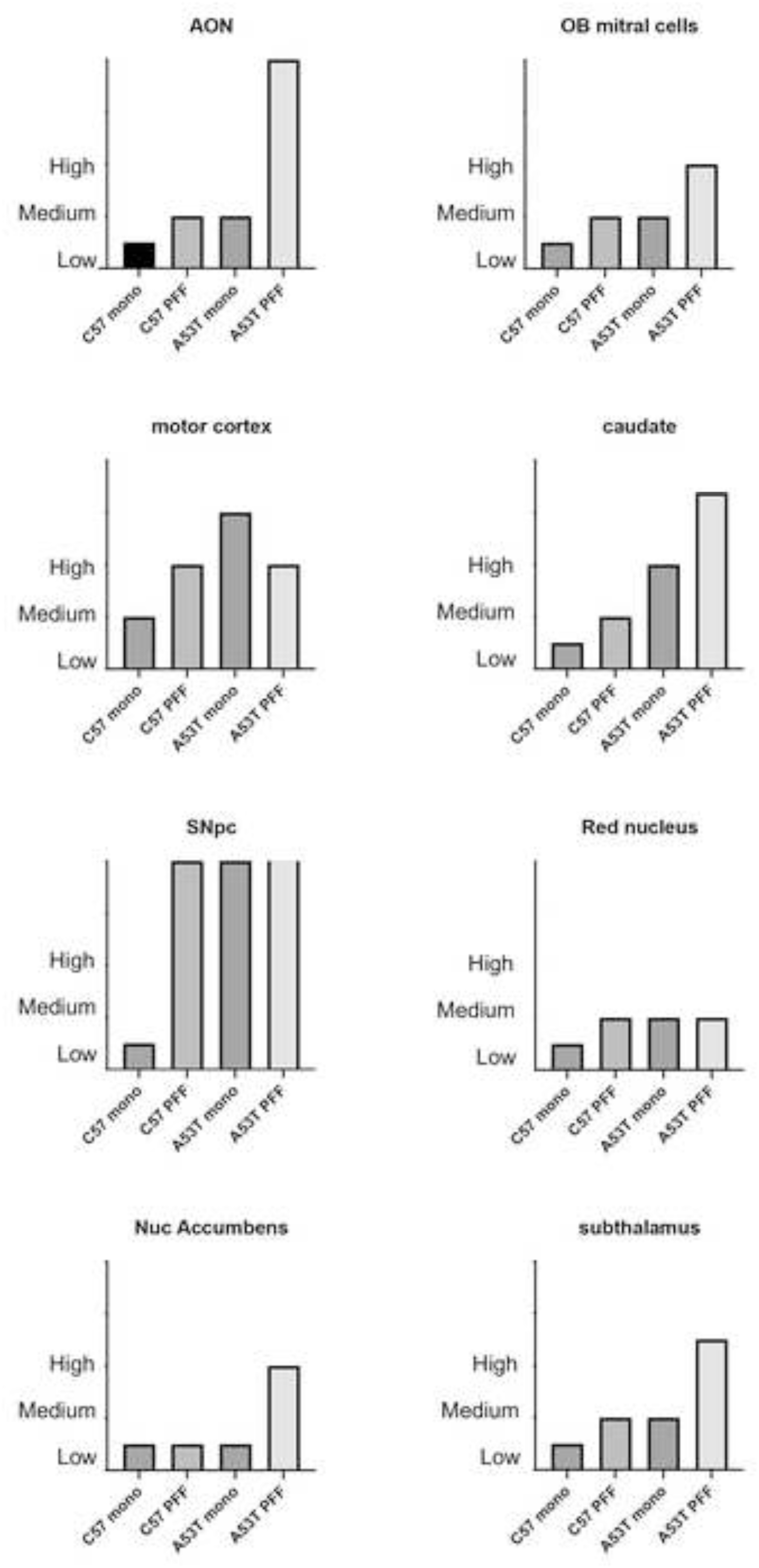
Relative burden of synucleinopathy following PFF injection into striatum. Different regions of the brain were examined for pSer129 α-syn burden. In all regions examined, injection of monomeric α-syn injection into the striatum of C57BL/6J mice resulted in low pSer129 α-syn burden (0-25% or 1^st^ quartile of animals showing any SYN transmission). PFF injection into the stratum of C57BL/6J mice resulted in low to moderate (25-50% or second quartile of mice showing transmission), except for the SNpc which showed high (>3^rd^ quartile) pSer129 α-syn burden. In A53T mice, injection of monomeric α-syn resulted in a synuclein burden scored as medium in all regions except the nuc. accumbens, except for the SNpc where pSer129 α-syn burden was 2 standard deviations above the average 4^th^ quartile score. When PFFs were injected into A53T striatum, pSer129 α-syn burden was scored as high (greater than 3^rd^ quartile in all regions examined, with AON, caudate, SNpc and subthalamus ranking at least 1 SD over the average 4^th^ quartile score.

To examine if the nigrostriatal pathway was at higher risk for pSer129 α-syn aggregation than other regions of the neuraxis, we injected a separate cohort of mice with PFFs into the rostral hippocampus. Like the striatal injections, we observed maximal transmission of insoluble pSer129 α-syn at 60 days, but the pattern of transmission was to structures that were primarily associated with the limbic system (**Suppl Fig 2**). In C57BL/6J mice that were injected with monomeric α-syn, in both striatum as well as hippocampus, we did not observe any pSer129 α-syn, with the exception of occasional aggregates directly at the injection sites (**Supple Table 1 and 2**).

In A53T mutant animals injected with PFFs in either the striatum or hippocampus, we observed the presence of higher levels of aggregates at each time point examined, with the same decrease in aggregation at 180dpi. Although higher levels of aggregation were seen in the A53T mice, the pattern of pSer129 α-syn was identical to that observed in the C57BL/6J mice. However, unlike the C57BL/6J mice, when monomeric α-syn was injected into the striatum or hippocampus of A53T mice, we observed pSer129 α-syn aggregates in the same pathways and at levels comparable to that seen after injection of PFFs (**Fig 2**). This suggests that the increased baseline levels of A53T α-syn protein in the transgenic mice (estimated at 2-3x protein compared to non-transgenic strain matched mice)[22] was sufficient to recruit the unfolded monomeric form of α-syn into the misfolded insoluble pSer129 α-syn. Based on this observation, one could infer that recruitment of monomeric forms of α-syn to misfolded oligomeric aggregates may be a mechanism for the finding of increased levels of insoluble α-syn in the brains of patients with duplication/triplication of the WT SNCA gene [29–33].

### Effect of PFF and monomeric α-Syn injection on SNpc DA neuron number

To further explore the effects of α-syn aggregation and transmission on SNpc health, we performed a stereologic evaluation of SNpc TH-positive DA neurons following monomeric and PFF injection to the striatum. In C57BL/6J mice, neither monomer nor PFF injection into the striatum (**Fig 3A,B,C**) resulted in any loss of SNpc DA neurons. However, in A53T mice, there was significant SNpc TH+ DA neuron loss at 60dpi and 180dpi in both PFF and monomer injected animals (**Fig 3A,B,D**). These data, paired with the results from the α-syn immunohistochemistry, were striking for two reasons. One, we did not see SNpc TH+ cell loss in the wild-type animals regardless of the form of α-syn injected. Secondly, we clearly demonstrate the ability of non-fibrillized α-syn monomer to both initiate aggregation and spread α-syn pathology, which we suggest induces a significant DA neuron loss in the SNpc. These results suggest that the presence of α-syn aggregates alone is not sufficient to drive neuron loss in the SNpc but may rely on the levels of induced neuroinflammation alone or in combination with α-syn burden.

**Fig. 3.**
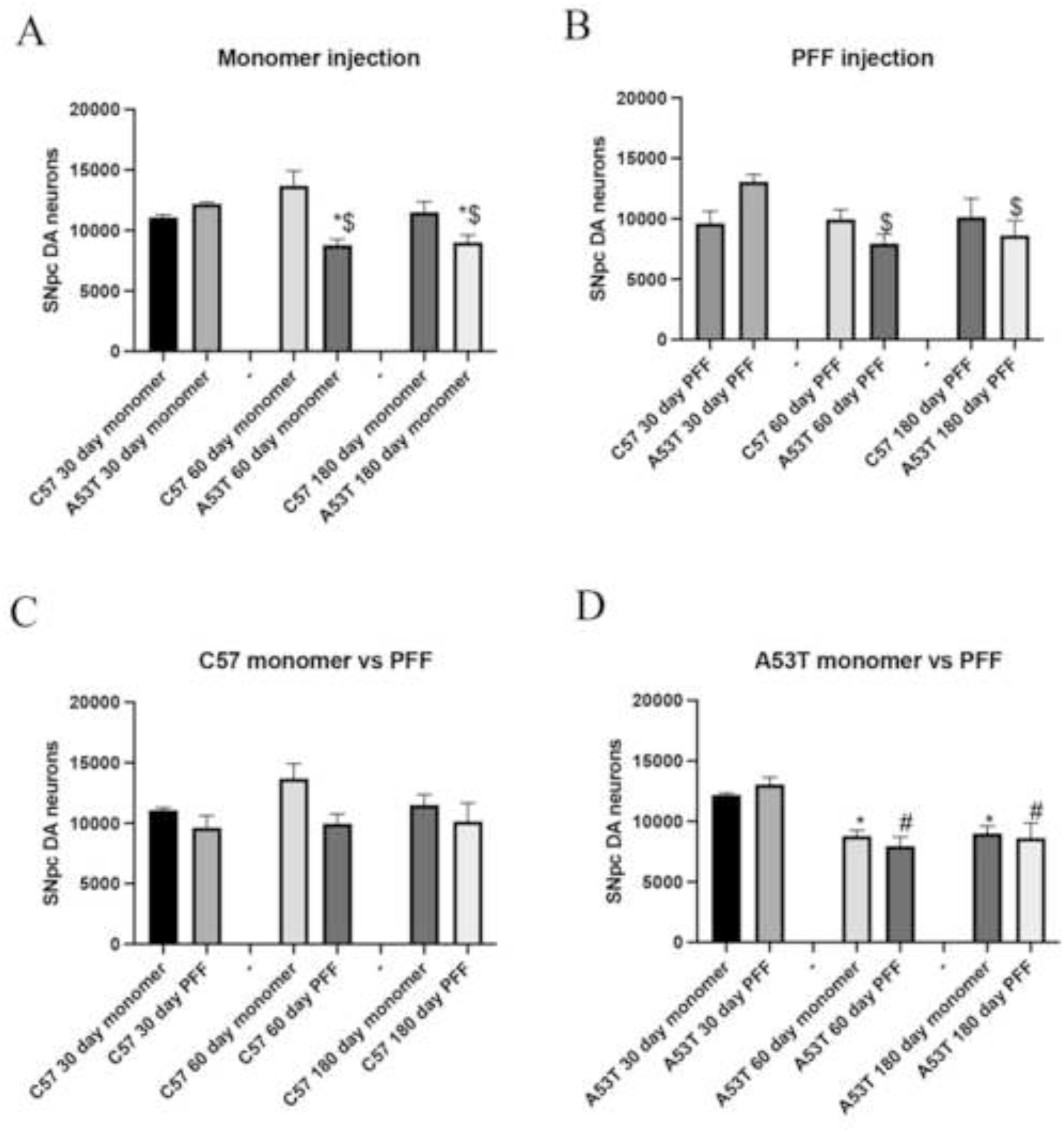
Effect of monomeric and PFF injection into striatum on SNpc DA neuron number. A). After monomeric α-syn injection, no loss of SNpc DA neurons were seen after 30, 60 or 180 days in C57BL/6J mice. However, in A53T mice, monomeric α-syn induced a significant loss of SNpc neurons at 60 and 180 days. B) After PFF injection, no loss of SNpc DA neurons were seen after 30, 60 or 180 days in C57BL/6J mice. However, in A53T mice, PFFs induced a significant loss of SNpc neurons at 60 and 180 days. * p<0.05 compared to same day monomer, $p<0.05 compared to 30 day timepoint C) Comparison of monomer versus PFFs in C57BL/6J mice. Neither monomeric α-syn nor PFFs induced a significant loss of SNpc neurons at 30, 60 or 180 days. D) Comparison of monomer versus PFFs in A53T mice. Neither monomeric α-syn nor PFFs induced a significant loss of SNpc neurons at 30 days post injection. However, at 60 and 180 days both the monomer and PFFs induced a significant and equal decrease in SNpc DA neuron number. * p<0.05 compared to 30 day monomer, # p<0.05 compared to 30 day PFF.

### Effect of PFF injection on Immune Response in C57BL/6J and A53T mice

To examine the immunological effects of intracerebral α-syn injection, we performed Iba-1 immunohistochemistry combined with stereologic techniques to assess how exogenous α-syn affected the number and state of SNpc microglia in the SNpc. While PFFs induced an increase in activated microglia when compared to monomer injections, we observed important differences dependent on the form of α-syn as well as the recipient animal’s genotype. In C57BL/6J mice, injection of monomeric forms of α-syn did not induce any significant inflammatory effect in the SNpc at any of the timepoints examined (**Fig 4A,B,C**). However, when PFFs were injected into dorsal striatum, and although extremely variable within groups, we measured an averaged 10-fold increase in the number of activated microglia at 30 dpi, which persisted through the 180 days of observation (**Fig 4B,C**). A quite different picture was seen in A53T mice. At 30 dpi, monomeric α-syn induced a similar microglial activation to that seen after PFFs in C57BL/6J mice (**Fig 4A**). The induction of microglia was similar in mice injected with PFFs at 30 dpi (**Fig 4A**). At 60 days, we observed a progressive loss of activated microglia in animals injected with monomeric α-syn, so that by 180 dpi, they had returned to baseline numbers (**Fig 4C**). In A53T mice injected with PFFs, the initial 30day increase in activated microglia was maintained for the entire 180-day period of observation.

**Fig 4.**
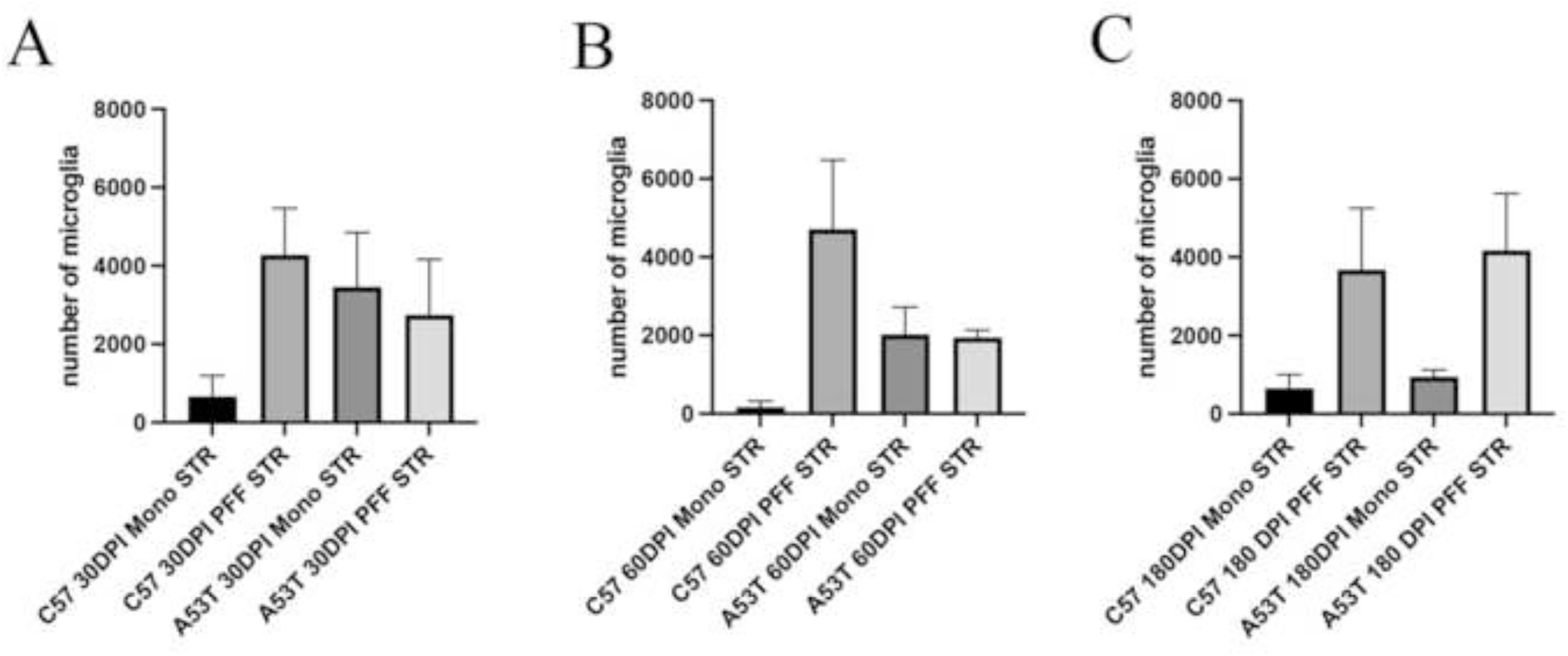
Stereological assessment of activated microglia in the SNpc following monomeric α-syn or PFF injection into striatum. A). 30 days after injection of monomeric or PFF α-syn we quantified a variable, but average 6-10 fold increase in the number of activated microglia in both C57BL/6J and A53T mice injected intrastriatally with PFFs. A similar increase was seen in A53T mice after monomeric α-syn. B) 60 days after injection of monomeric or PFF α-syn we quantified a variable, but average 6-10 fold increase in the number of activated microglia in both C57BL/6J and A53T mice injected intrastriatally with PFFs. C) 180 days after injection of monomeric of PFF α-syn we quantified a variable, but average 8-10 fold increase in the number of activated microglia only in mice injected intrastriatally with PFFs. The number of activated microglia in A53T injected with monomeric mice α-syn was similar to that seen in C57BL/6J mice.

We also examined the expression of 4 proinflammatory cytokines (IFNg, TNFα, IL-6 and IL-1a) and one anti-inflammatory cytokine (IL-10) in the SNpc following intrastriatal injection of PFFs. In C57BL/6J mice, we measured a significant increase in each of the proinflammatory cytokines 30 days after PFF injection. By 60 dpi, IFNg, TNFα, and IL-1a had returned to baseline levels, while IL-6 levels remained elevated (**Fig 5A,C,E,G**). Like the C57BL/6J mice, striatal PFF injection into A53T mice resulted in a significant induction of all 4 proinflammatory cytokines 30 dpi. However, unlike the C57BL/6J animals, the A53T mice demonstrated a continued elevation of these inflammatory mediators through the entire experimental period (180 dpi) (**Fig. 5B,D,F,H**). We also examined changes in the anti-inflammatory cytokine IL-10. C57BL/6J mice had significant increases at 30 and 60 dpi, which returned to baseline levels at 18o dpi (**Fig 5I**). In A53T mice, we also measured a significant increase in IL-10 (**Fig 5J**) but its induction compared to the C57BL/6J was only reduced by approximately 65% (**Fig 5I,J**).

**Fig 5.**
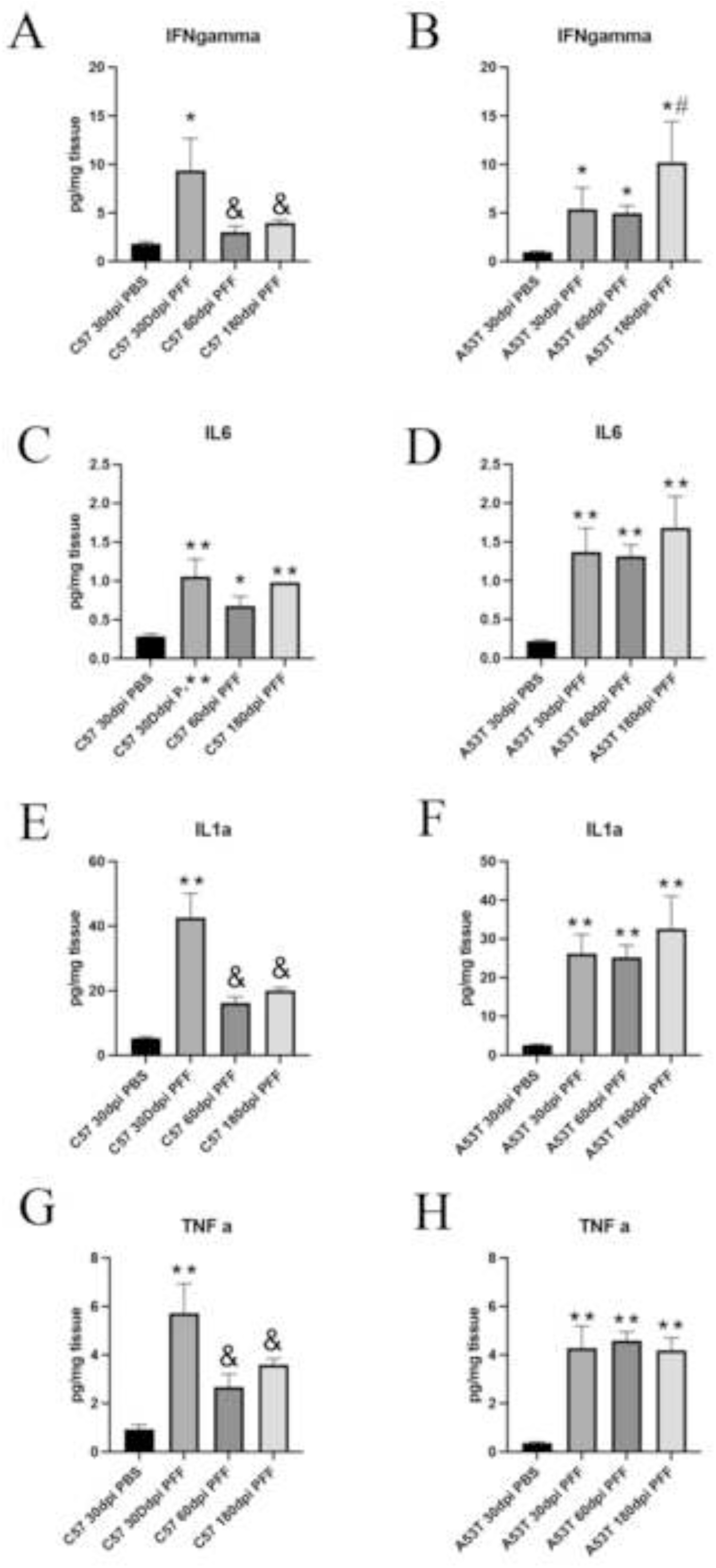
SNpc cytokine activation 30 days following intrastriatal injection of PFFs in C57BL/6J or A53T SNCA mice. A) Expression of IFNγ was significantly increased 30 days after injection of PFFs in C57BL/6J mice compared to mice injected with PBS. At both 60 and 180 days after injection of PFFs, we see a significant increase in both C57BL/6J and mice compared to saline injected mice, but at this time, these levels have significantly been reduced compared to 30 days after PFFs. B) Expression of IFNγ was significantly increased 30, 60 and 180 days after injection of PFFs in A53T SNCA mice compared to A53T SNCA mice injected with PBS. We also measured a significant increase in IFNγ in AA53T SNCA mice compared to levels measured at 30 or 60 days after PFF injection. C) Expression of IL-6 was significantly increased 30, 60 and 180 days after injection of PFFs in C57BL/6J mice compared to C57BL/6J mice injected with PBS. D) Expression of IL-6 was significantly increased 30, 60 and 180 days after injection of PFFs in A53T SNCA mice compared to mice injected with PBS. E) Expression of IL1a was significantly increased 30, 60 and 180 days after injection of PFFs in C57BL/6J mice compared to C57BL/6J mice injected with PBS. F) Expression of IL1a was significantly increased 30, 60 and 180 days after injection of PFFs in A53T SNCA mice compared to A53T SNCA mice injected with PBS. G) Expression of TNFa was significantly increased 30, 60 and 180 days after injection of PFFs in C57BL/6J mice compared to C57BL/6J mice injected with PBS but at this time, TNFa levels have significantly been reduced compared to 30 days after PFFs. H) Expression of TNFa was significantly increased 30, 60 and 180 days after injection of PFFs in A53T SNCA mice compared to A53T SNCA mice injected with PBS. I) Expression of IL-10 was significantly increased 30, 60 and 180 days after injection of PFFs in C57BL/6J mice compared to C57BL/6J mice injected with PBS. J) Expression of IL-10 was significantly increased at 60dpi and 180dpi although it was 65% less than C57BL/6J mice. ** p<0.01 compared to saline, *p<0.05 compared to saline. & p<0.05 compared to 30 day.

Given the variability within groups of the number of activated microglia induced by monomer of PFF injection, we performed a regression analysis of individual animals to correlate the number of activated microglia with SNpc DA neuron loss. As seen in **Fig 6**, we did not detect any significant correlation between these two variables. This suggested that the cell death seen after PFF injection in both C57BL/6J and A53T mice was likely due to direct synuclein burden. The direct role of inflammation as an inducer of neuron death, based on our results, is controversial. While we did not find any correlation between the number of activated microglia/animal and SNpc DA neuron death, we do find significant increases in proinflammatory cytokines following injection of PFFs. This suggests that it is these chemical mediators rather than the number of activated microglia themselves that serve as a marker of the degeneration observed.

**Fig 6.**
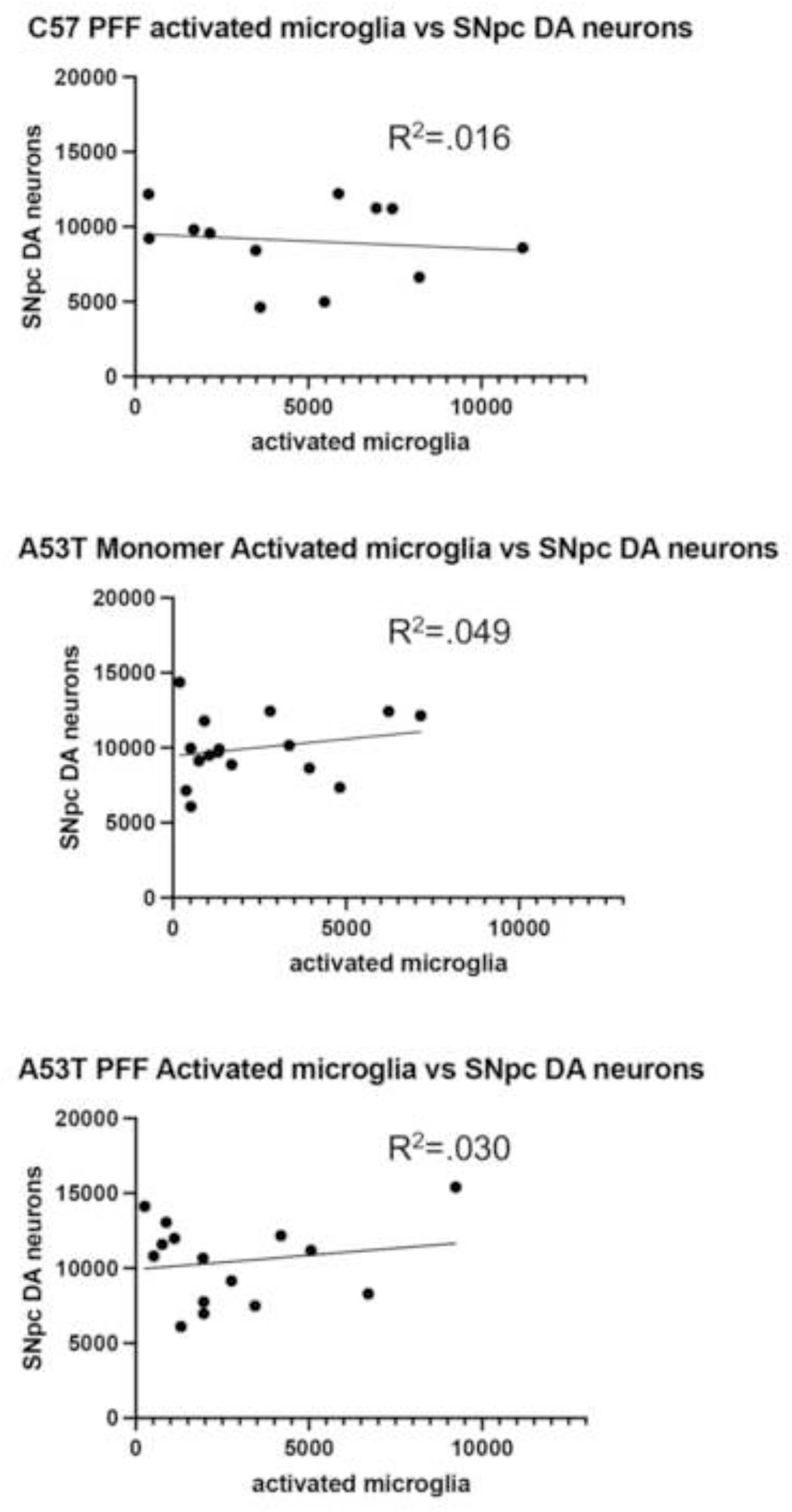
Correlation between activated microglia and SNpc DA neuron loss. No positive or negative correlation was measured in C57BL/6J PFF injected or A53T SNCA mice after monomeric or PFF injection into striatum. Each dot represents data from 1 animal.

## Discussion

Our findings in this study, using a PFF model of induced synucleinopathy, support previous work that has demonstrated the potential for specific transmission of oligomeric α-syn fragments [18, 34–36]. We also show that the transmission of these fragments is not only permissive in the nigrostriatal tract, but can also occur in other synaptically-connected regions. Additionally, we also show that monomeric forms of α-syn can also be recruited into insoluble aggregates in brains when the endogenous α-syn burden is already at higher than physiological levels. This observation may provide insight that provides a mechanism for the early onset of parkinsonism seen in humans that carry duplication/triplication, without any mutation, of the SNCA gene [31, 37].

While the presence of insoluble α-syn inclusions in the SNpc- and throughout the neuroaxis-are hallmarks of PD, there is active debate about the correlation between α-syn burden and degree of neuronal dysfunction. Studies in animals offer evidence that formation of inclusions coincides with neuronal dysfunction, while others show a weak correlation between inclusion burden and loss of SNpc TH neurons after striatal PFF injections in rodent models. [38, 39]. However, in post-mortem tissue of PD patients, the relationship between protein load and SN degeneration is unclear, with some patients experiencing PD symptoms and loss of nigral neurons without α-syn aggregates.[40, 41] Additionally, Lewy pathology is present in other neurodegenerative diseases that are clinically different from PD [42]. While the relationship between the spread of Lewy pathology and clinical progression of PD has been outlined by Braak and his colleagues [13, 43], only about half of people with clinical PD have a distribution of Lewy pathology that exhibits the classic pattern of Braak staging, and those with genetic forms of disease can be even more distinct from this pattern of synuclein deposition [44]. Critical to our understanding of α-syn spread is the finding that Lewy pathology is hypothesized to spread through synaptically-connected networks of neurons, rather than through its nearest neighbor [19, 20]. While this hypothesis has clear support from our injections to striatum (progression in the nigrostriatal system) and hippocampus (progression in the limbic system), there are reports that some regions of high Lewy pathology burden, such as locus coeruleus, do not act as nodes of propagation [45, 46]. The heterogeneity in PD and related synucleinopathies between Lewy body formation and neurodegeneration suggests there are distinct regional brain differences that could be governing this relationship. It has been shown that there are differences within the midbrain (medial vs. ventrolateral SN, and SN vs. VTA) regarding axonal arbor dimension, pacemaking mechanisms, calcium signaling, and mitochondrial oxidative stress that results in various levels of cell vulnerability in neurodegeneration.[47]. With such variability in the effects of Lewy pathology in various brain regions, we focused on characterizing inflammation which could be a crucial factor in understanding the cellular context of neurodegeneration in synucleinopathies.

Regarding our examination of inflammation, we found no correlation between the number of activated microglia in the SNpc and the loss of SNpc DA neurons. However, we did find that when SNpc DA neurons were lost there was a significant increase in the expression of proinflammatory cytokines in this tissue. This experimental finding is supported by the observation, in humans, of microglia that immunostain for the pro-inflammatory cytokines TNFα and IL-6 [48]. This suggests, at least following induction from overexpression of α-syn, that the morphological appearance of these inflammatory cells cannot, alone, be used as a marker of toxic inflammation, but should be used in conjunction with direct measurement of their inflammatory secretome. Additionally, while there is no conclusive evidence from our studies, it does seem that the increase in proinflammatory cytokines does precede SNpc DA neuron death. This suggests to us that inflammation along with increased α-syn burden is required for SNpc DA neuron death to occur [49–52]. The role of PFFs in driving this inflammation may also be related to the brain’s overall synuclein burden. For example, we do not see any rise in microglial number-nor cytokine expression-in C57BL/6J mice injected with monomeric forms of α-syn; but do see this in A53T mice. While it has been shown that the mutant A53T form of α-syn is capable of more strongly activating wild-type microglia in mice, it was previously unknown how mutant A53T microglia are able to respond to exogenous wild-type human α-syn [51]. While we do not know if the form of synuclein drives inflammation or if the protein must first template into oligomeric forms, it has been reported that α-syn monomer does not activate microglia in vitro [53] The mutant mice we used in this study overexpressed human A53T synuclein, resulting in a 3.2 fold overexpression of α-syn on a mouse SNCA null background; thus all α-syn is human in form. It is possible that this overexpression of soluble α-synuclein puts stress on the microglial load for handling protein degradation, such that exogenous application induced an overall pro-inflammatory response. If this is the case, then two possible deleterious consequences could arise: 1) microglia have reached maximum capacity for protein degradation and cannot maintain protein homeostasis. Mechanistically, this would occur via a gain of inflammatory phenotype or loss of anti-inflammatory phenotype, furthering cellular stress and protein propagation, or 2) the microglial activity is altered, potentially leading to the development of more toxic species of α-syn oligomers that subsequently drive propagation and inflammation. Additionally, we know that monomeric α-syn is capable of interacting with aggregated α-syn via intramolecular unfolding, which promotes further protein aggregation.[54] While it has not been previously reported in the A53T double PAC mutant mice, M83 A53T mice do report increasing levels of oligomeric α-syn with age, with the A53T double PAC mice having some evidence of aggregated α-syn in the ENS and dystrophic neurites in the hippocampus [55, 56] Since there may be increased levels of aggregated α-syn in the A53T double PAC mice, it is possible that intrastriatal monomeric injections of α-syn are capable of directly interacting with oligomeric species to accelerate pathology. Furthermore, not only is the proteomic environment permissive of aggregation, but also the transgenic immune environment, namely microglia, could contribute to the aggregation process.

For this reason, the handling and processing of α-syn by microglia must be central to future studies. Our results reinforce this pivotal role that microglia play in homeostasis of the nigrostriatal pathway. Here, we demonstrate that in mice overexpressing A53T α-syn, that any exogenous application of α-syn, monomeric or fibrillized, is capable of inducing an inflammatory cascade and suppressing anti-inflammatory signaling. We also find that the inflammatory secretome play a central role both in the response to α-syn aggregation as well as in mediating pro- and anti-inflammatory responses in a PFF model of synucleinopathy. In terms of disease progression, our finding of monomeric induction of a parkinsonian pathology in mice that overexpress the A53T form of this mutation-which is the WT form in mice [57] may provide a mechanism to explain the earlier onset of disease in persons carrying duplication/triplication of the WT SNCA gene.

## Figure Legends

**<u>Suppl Fig 1. Map of pSer-129 aggregation in C57BL/6J and A53T mice following injection of PFFs into the rostral hippocampus</u>.** A. 60 days after injection of PFFs into the CA3 region of the rostral hippocampus we observed (from rostral to caudal) pSer129 α-syn immunoreactive aggregation throughout the neuraxis. Areas highlighted in yellow showed low to medium pSer129 α-syn aggregate burden. These structures are limbic in nature. AON: accessory olfactory nucleus.

## Acknowledgements

The Neurochemistry Core is supported by the Vanderbilt Brain Institute and the Vanderbilt Kennedy Center. The work in this manuscript was supported in part by funding by the Michael J Fox Foundation for Parkinson’s Research. Dr. Smeyne is an Editorial Board Member of this journal, but was not involved in the peer-review process nor had access to any information regarding its peer-review.’’

## Notes

### Competing Interest Statement

The authors have declared no competing interest.

## References

[1] (2019) Global, regional, and national burden of neurological disorders, 1990-2016: a systematic analysis for the Global Burden of Disease Study 2016. Lancet Neurol 18, 459–480.

[2] Steece-Collier K, Maries E, Kordower JH (2002) Etiology of Parkinson’s disease: Genetics and environment revisited. Proceedings of the National Academy of Sciences 99, 13972–13974.

[3] Chaudhuri KR, Healy DG, Schapira AH (2006) Non-motor symptoms of Parkinson’s disease: diagnosis and management. Lancet Neurol 5, 235–245.

[4] Bloem BR, Okun MS, Klein C (2021) Parkinson’s disease. Lancet 397, 2284–2303.

[5] Mari Z, Mestre TA (2022) The Disease Modification Conundrum in Parkinson’s Disease: Failures and Hopes. Front Aging Neurosci 14, 810860.

[6] Chu Y, Hirst WD, Kordower JH (2023) Mixed pathology as a rule, not exception: Time to reconsider disease nosology. Handb Clin Neurol 192, 57–71.

[7] Weiss F, Labrador-Garrido A, Dzamko N, Halliday G (2022) Immune responses in the Parkinson’s disease brain. Neurobiol Dis 168, 105700.

[8] Kalia LV, Lang AE (2015) Parkinson’s disease. Lancet 386, 896–912.

[9] Polymeropoulos MH, Lavedan C, Leroy E, Ide SE, Dehejia A, Dutra A, Pike B, Root H, Rubenstein J, Boyer R, Stenroos ES, Chandrasekharappa S, Athanassiadou A, Papapetropoulos T, Johnson WG, Lazzarini AM, Duvoisin RC, Di Iorio G, Golbe LI, Nussbaum RL (1997) Mutation in the alpha-synuclein gene identified in families with Parkinson’s disease. Science 276, 2045–2047.

[10] Simon-Sanchez J, Schulte C, Bras JM, Sharma M, Gibbs JR, Berg D, Paisan-Ruiz C, Lichtner P, Scholz SW, Hernandez DG, Kruger R, Federoff M, Klein C, Goate A, Perlmutter J, Bonin M, Nalls MA, Illig T, Gieger C, Houlden H, Steffens M, Okun MS, Racette BA, Cookson MR, Foote KD, Fernandez HH, Traynor BJ, Schreiber S, Arepalli S, Zonozi R, Gwinn K, van der Brug M, Lopez G, Chanock SJ, Schatzkin A, Park Y, Hollenbeck A, Gao J, Huang X, Wood NW, Lorenz D, Deuschl G, Chen H, Riess O, Hardy JA, Singleton AB, Gasser T (2009) Genome-wide association study reveals genetic risk underlying Parkinson’s disease. Nat Genet 41, 1308–1312.

[11] Jellinger KA (2014) The pathomechanisms underlying Parkinson’s disease. Expert Rev Neurother 14, 199–215.

[12] Bousset L, Pieri L, Ruiz-Arlandis G, Gath J, Jensen PH, Habenstein B, Madiona K, Olieric V, Böckmann A, Meier BH, Melki R (2013) Structural and functional characterization of two alpha-synuclein strains. Nat Commun 4, 2575.

[13] Braak H, Del Tredici K, Rub U, de Vos RA, Jansen Steur EN, Braak E (2003) Staging of brain pathology related to sporadic Parkinson’s disease. Neurobiol Aging 24, 197–211.

[14] Kordower JH, Brundin P (2009) Lewy body pathology in long-term fetal nigral transplants: is parkinson’s disease transmitted from one neural system to another? Neuropsychopharmacology 34, 254–254.

[15] Heneka MT, Kummer MP, Latz E (2014) Innate immune activation in neurodegenerative disease. Nat Rev Immunol 14, 463–477.

[16] Zhang W, Wang T, Pei Z, Miller DS, Wu X, Block ML, Wilson B, Zhang W, Zhou Y, Hong JS, Zhang J (2005) Aggregated alpha-synuclein activates microglia: a process leading to disease progression in Parkinson’s disease. Faseb j 19, 533–542.

[17] Moore DJ, West AB, Dawson VL, Dawson TM (2005) MOLECULAR PATHOPHYSIOLOGY OF PARKINSON’S DISEASE. Annual Review of Neuroscience 28, 57–87.

[18] Luk KC, Kehm V, Carroll J, Zhang B, O’Brien P, Trojanowski JQ, Lee VM (2012) Pathological alpha-synuclein transmission initiates Parkinson-like neurodegeneration in nontransgenic mice. Science 338, 949–953.

[19] Luk KC, Kehm VM, Zhang B, O’Brien P, Trojanowski JQ, Lee VM (2012) Intracerebral inoculation of pathological α-synuclein initiates a rapidly progressive neurodegenerative α-synucleinopathy in mice. J Exp Med 209, 975–986.

[20] Volpicelli-Daley LA, Luk KC, Patel TP, Tanik SA, Riddle DM, Stieber A, Meaney DF, Trojanowski JQ, Lee VM (2011) Exogenous α-synuclein fibrils induce Lewy body pathology leading to synaptic dysfunction and neuron death. Neuron 72, 57–71.

[21] Rey NL, Steiner JA, Maroof N, Luk KC, Madaj Z, Trojanowski JQ, Lee VM, Brundin P (2016) Widespread transneuronal propagation of α-synucleinopathy triggered in olfactory bulb mimics prodromal Parkinson’s disease. J Exp Med 213, 1759–1778.

[22] Kuo YM, Li Z, Jiao Y, Gaborit N, Pani AK, Orrison BM, Bruneau BG, Giasson BI, Smeyne RJ, Gershon MD, Nussbaum RL (2010) Extensive enteric nervous system abnormalities in mice transgenic for artificial chromosomes containing Parkinson disease-associated {alpha}-synuclein gene mutations precede central nervous system changes. Hum Mol Genet.

[23] Baquet ZC, Williams D, Brody J, Smeyne RJ (2009) A comparison of model-based (2D) and design-based (3D) stereological methods for estimating cell number in the substantia nigra pars compacta (SNpc) of the C57BL/6J Mouse. Neuroscience 161, 1082–1090.

[24] Sadasivan S, Zanin M, O’Brien K, Schultz-Cherry S, Smeyne RJ (2015) Induction of microglia activation after infection with the non-neurotropic A/CA/04/2009 H1N1 influenza virus. PLoS One 10, e0124047.

[25] Graeber MB, Streit WJ (2010) Microglia: biology and pathology. Acta Neuropathol 119, 89–105.

[26] Paxinos G, Franklin KBJ (2001) The Mouse Brain in Stereotaxic Coordinates, Academic Press, San Diego.

[27] Chen SC, Ehrhard P, Goldowitz D, Smeyne RJ (1997) Developmental expression of the GIRK family of inward rectifying potassium channels: implications for abnormalities in the weaver mutant mouse. Brain Res 778, 251–264.

[28] Chen SC, Kochan JP, Campfield LA, Burn P, Smeyne RJ (1999) Splice variants of the OB receptor gene are differentially expressed in brain and peripheral tissues of mice. J Recept Signal Transduct Res 19, 245–266.

[29] Becerra-Calixto A, Mukherjee A, Ramirez S, Sepulveda S, Sinha T, Al-Lahham R, De Gregorio N, Gherardelli C, Soto C (2023) Lewy Body-like Pathology and Loss of Dopaminergic Neurons in Midbrain Organoids Derived from Familial Parkinson’s Disease Patient. Cells 12.

[30] Chartier-Harlin MC, Kachergus J, Roumier C, Mouroux V, Douay X, Lincoln S, Levecque C, Larvor L, Andrieux J, Hulihan M, Waucquier N, Defebvre L, Amouyel P, Farrer M, Destée A (2004) Alpha-synuclein locus duplication as a cause of familial Parkinson’s disease. Lancet 364, 1167–1169.

[31] Konno T, Ross OA, Puschmann A, Dickson DW, Wszolek ZK (2016) Autosomal dominant Parkinson’s disease caused by SNCA duplications. Parkinsonism Relat Disord 22 Suppl 1, S1–6.

[32] Olgiati S, Thomas A, Quadri M, Breedveld GJ, Graafland J, Eussen H, Douben H, de Klein A, Onofrj M, Bonifati V (2015) Early-onset parkinsonism caused by alpha-synuclein gene triplication: Clinical and genetic findings in a novel family. Parkinsonism Relat Disord 21, 981–986.

[33] Zafar F, Valappil RA, Kim S, Johansen KK, Chang ALS, Tetrud JW, Eis PS, Hatchwell E, Langston JW, Dickson DW, Schüle B (2018) Genetic fine-mapping of the Iowan SNCA gene triplication in a patient with Parkinson’s disease. NPJ Parkinsons Dis 4, 18.

[34] Paumier KL, Luk KC, Manfredsson FP, Kanaan NM, Lipton JW, Collier TJ, Steece-Collier K, Kemp CJ, Celano S, Schulz E, Sandoval IM, Fleming S, Dirr E, Polinski NK, Trojanowski JQ, Lee VM, Sortwell CE (2015) Intrastriatal injection of pre-formed mouse alpha-synuclein fibrils into rats triggers alpha-synuclein pathology and bilateral nigrostriatal degeneration. Neurobiol Dis 82, 185–199.

[35] Volpicelli-Daley LA, Luk KC, Patel TP, Tanik SA, Riddle DM, Stieber A, Meaney DF, Trojanowski JQ, Lee VM (2011) Exogenous alpha-synuclein fibrils induce Lewy body pathology leading to synaptic dysfunction and neuron death. Neuron 72, 57–71.

[36] Kim S, Kwon SH, Kam TI, Panicker N, Karuppagounder SS, Lee S, Lee JH, Kim WR, Kook M, Foss CA, Shen C, Lee H, Kulkarni S, Pasricha PJ, Lee G, Pomper MG, Dawson VL, Dawson TM, Ko HS (2019) Transneuronal Propagation of Pathologic α-Synuclein from the Gut to the Brain Models Parkinson’s Disease. Neuron 103, 627–641.e627.

[37] Olanow CW, Brundin P (2013) Parkinson’s disease and alpha synuclein: is Parkinson’s disease a prion-like disorder? Mov Disord 28, 31–40.

[38] Osterberg VR, Spinelli KJ, Weston LJ, Luk KC, Woltjer RL, Unni VK (2015) Progressive aggregation of alpha-synuclein and selective degeneration of lewy inclusion-bearing neurons in a mouse model of parkinsonism. Cell Rep 10, 1252–1260.

[39] Abdelmotilib H, Maltbie T, Delic V, Liu Z, Hu X, Fraser KB, Moehle MS, Stoyka L, Anabtawi N, Krendelchtchikova V, Volpicelli-Daley LA, West A (2017) alpha-Synuclein fibril-induced inclusion spread in rats and mice correlates with dopaminergic Neurodegeneration. Neurobiol Dis 105, 84–98.

[40] Jellinger KA (2009) A critical evaluation of current staging of alpha-synuclein pathology in Lewy body disorders. Biochim Biophys Acta 1792, 730–740.

[41] Surmeier DJ, Obeso JA, Halliday GM (2017) Selective neuronal vulnerability in Parkinson disease. Nat Rev Neurosci 18, 101–113.

[42] Dijkstra AA, Voorn P, Berendse HW, Groenewegen HJ, Rozemuller AJ, van de Berg WD (2014) Stage-dependent nigral neuronal loss in incidental Lewy body and Parkinson’s disease. Mov Disord 29, 1244–1251.

[43] Del Tredici K, Braak H (2016) Review: Sporadic Parkinson’s disease: development and distribution of α-synuclein pathology. Neuropathol Appl Neurobiol 42, 33–50.

[44] Halliday G, McCann H, Shepherd C (2012) Evaluation of the Braak hypothesis: how far can it explain the pathogenesis of Parkinson’s disease? Expert Rev Neurother 12, 673–686.

[45] Schwarz LA, Miyamichi K, Gao XJ, Beier KT, Weissbourd B, DeLoach KE, Ren J, Ibanes S, Malenka RC, Kremer EJ, Luo L (2015) Viral-genetic tracing of the input-output organization of a central noradrenaline circuit. Nature 524, 88–92.

[46] Halliday GM, Song YJ, Harding AJ (2011) Striatal β-amyloid in dementia with Lewy bodies but not Parkinson’s disease. J Neural Transm (Vienna) 118, 713–719.

[47] Liss B, Roeper J (2008) Individual dopamine midbrain neurons: functional diversity and flexibility in health and disease. Brain Res Rev 58, 314–321.

[48] Imamura K, Hishikawa N, Sawada M, Nagatsu T, Yoshida M, Hashizume Y (2003) Distribution of major histocompatibility complex class II-positive microglia and cytokine profile of Parkinson’s disease brains. Acta Neuropathol 106, 518–526.

[49] Anderson FL, Biggs KE, Rankin BE, Havrda MC (2023) NLRP3 inflammasome in neurodegenerative disease. Transl Res 252, 21–33.

[50] Grotemeyer A, Fischer JF, Koprich JB, Brotchie JM, Blum R, Volkmann J, Ip CW (2023) Inflammasome inhibition protects dopaminergic neurons from α-synuclein pathology in a model of progressive Parkinson’s disease. J Neuroinflammation 20, 79.

[51] Hoenen C, Gustin A, Birck C, Kirchmeyer M, Beaume N, Felten P, Grandbarbe L, Heuschling P, Heurtaux T (2016) Alpha-Synuclein Proteins Promote Pro-Inflammatory Cascades in Microglia: Stronger Effects of the A53T Mutant. PLoS One 11, e0162717.

[52] Leem YH, Kim DY, Park JE, Kim HS (2023) Necrosulfonamide exerts neuroprotective effect by inhibiting necroptosis, neuroinflammation, and α-synuclein oligomerization in a subacute MPTP mouse model of Parkinson’s disease. Sci Rep 13, 8783.

[53] Li N, Stewart T, Sheng L, Shi M, Cilento EM, Wu Y, Hong J-S, Zhang J (2020) Immunoregulation of microglial polarization: an unrecognized physiological function of α-synuclein. Journal of Neuroinflammation 17, 272.

[54] Kumari P, Ghosh D, Vanas A, Fleischmann Y, Wiegand T, Jeschke G, Riek R, Eichmann C (2021) Structural insights into α-synuclein monomer-fibril interactions. Proc Natl Acad Sci U S A 118.

[55] Kuo YM, Li Z, Jiao Y, Gaborit N, Pani AK, Orrison BM, Bruneau BG, Giasson BI, Smeyne RJ, Gershon MD, Nussbaum RL (2010) Extensive enteric nervous system abnormalities in mice transgenic for artificial chromosomes containing Parkinson disease-associated alpha-synuclein gene mutations precede central nervous system changes. Hum Mol Genet 19, 1633–1650.

[56] Tsika E, Moysidou M, Guo J, Cushman M, Gannon P, Sandaltzopoulos R, Giasson BI, Krainc D, Ischiropoulos H, Mazzulli JR (2010) Distinct region-specific alpha-synuclein oligomers in A53T transgenic mice: implications for neurodegeneration. J Neurosci 30, 3409–3418.

[57] Hong L, Ko HW, Gwag BJ, Joe E, Lee S, Kim YT, Suh YH (1998) The cDNA cloning and ontogeny of mouse alpha-synuclein. Neuroreport 9, 1239–1243.

