## Supplemental table 1 for "Templating of Monomeric Alpha-Synuclein Induces Inflammation and SNpc Dopamine Neuron Death in a Genetic Mouse Model of Synucleinopathy"

**Supplemental Table 1**
**Expression of pSer129 in brain after injections**

| **Olfactory Bulb** | cells | **Diencephalon and Subcortical Areas** | cells | **Midbrain and Brainstem** | cells | |
|---|---|---|---|---|---|---|
| mitral cells | | DM hypothalamus | | vent. tegmental area | | Brain: |
| granule cells | | Dorsal endopiriform | | raphe nucleus | | |
| anterior olf. nuc. | | VM hypothalamus | | central grey | | |
| internal plexiform cells | | mammillary nucleus | | MGN | | Condition |
| periglomerular cells | | habenular nuc. | | LGN | | |
| **Cerebral Cortices and Telencephalon** | | subfornical organ | | D. tegmental nuc. | | |
| Piriform | | SCN | | med. vestib nuc. | | |
| Ant. Cingulate | | Lateral hypothal | | lat. Vestib nuc. | | Date Analyzed |
| Motor | | MGN | | pontine nuc. | | |
| Sensory | | preoptic area | | solitary nuc. | | |
| Parietal cortex | | zona incerta | | Cuneiform nuc. | | |
| Entorhinal cortex | | subiculum | | Periolivary n | | |
| Frontal cortex | | reuniens nuc. | | Reticular nuc. | | |
| Infralimbic cortex | | M. cerebellar ped. | | Pedunculopontine | | |
| Orbital cortex | | paraventricular hypothalamic nuc. | | Deep mesencephalic n | | |
| caudatoputamen | | BN stria terminalis | | **Cerebellum** | | |
| **Diencephalon and Subcortical Areas** | | median eminence | | EGL | | |
| Amygdala | | **Hippocampus** | | IGL | | |
| Tenia tecta | | CA1 | | Purkinje cells | | |
| Lateral Septal nuc. | | CA2 | | Nuclear cells | | |
| Med. Septal nuc. | | CA3 | | Molecular layer | | |
| Ventral pallidum | | CA4 | | choroid plexus | | |
| Indusium Griseum | | dentate gyrus | | **Cranial nerves** | | |
| Isles of Calleja | | **Midbrain and Brainstem** | | Nuc. III | | |
| Nuc. accumbens | | IPN | | Nuc. IV | | |
| Ant. tier thalamus | | area postrema | | Nuc. V | | |
| Vent. tier thalamus | | locus coeruleus | | MLF | | |
| Lat. tier thalamus | | Substantia nigra ret | | Post commissure | | |
| parafascicular nuc. | | substantia nigra, compacta | | Lateral lemniscus | | |
| Post. thalamic nuc. | | red nucleus | | Cerebral peduncle | | |
| Ant. hypothalamus | | Subthalamic nuc | | Corpus callosum | | |
| post. hypothalamus | | sup. colliculus | | Anterior commissure | | |
| reticular thalamus | | Inf colliculus | | fornix | | |
| arcuate hypothal. | | A8 retrorubral field | | Internal capsule | | |
| paraventricular thal | | sup. mammillary nuc. | | | | |
